# Nonlinear Relationship Between the C-Reactive Protein to Albumin Ratio and Chronic Kidney Disease: A Cross-Sectional Analysis Using the NHANES Database

**DOI:** 10.1101/2025.02.07.637012

**Authors:** Quanyuan Huang, Xiao Lu, Yuanhua Liang, Wuchang Zhu, Jinglan Chen, Yufang Yang

**Author notes:** Correspondence: Yufang Yang.

## Abstract

**Objective:** The C-reactive protein to albumin ratio (CAR) has gained attention as an inflammatory biomarker with potential relevance in various diseases. However, its relationship with chronic kidney disease (CKD) remains inadequately understood. This study aims to investigate the association between CAR and CKD prevalence in a diverse, multi-ethnic U.S. cohort.

**Methods:** This cross-sectional analysis utilized data from the 1999–2010 National Health and Nutrition Examination Survey (NHANES). Multivariable regression and subgroup analyses were conducted to control for confounding factors and examine the link between CAR and CKD. A restrictive cubic spline model was employed to explore potential nonlinear associations.

**Results:** Among 30,049 participants aged 18 years or older, 5,423 (18.05%) had CKD. Compared to the lowest quartile (Q1) of log-transformed CAR, CKD risk was higher in the second (Q2), third (Q3), and fourth (Q4) quartiles, with odds ratios of 1.28, 1.29, and 1.63, respectively. A dose-response relationship was observed, indicating that higher CAR levels were significantly associated with increased CKD risk.

**Conclusion:** Elevated CAR is independently associated with a higher prevalence of CKD, suggesting its potential utility as a biomarker for CKD risk assessment in diverse populations. Further studies are needed to confirm these findings and explore underlying mechanisms.

## 1 Introduction

Chronic Kidney Disease (CKD) is a progressive condition marked by sustained kidney damage or functional decline lasting over three months. Pathologically, CKD is characterized by significant injury to the renal tubules and glomeruli, resulting in tubular dysfunction and reduced glomerular filtration rate[1]. With the increasing global burden of chronic diseases, CKD has become a major public health challenge. The pathogenesis of this disease is complex, involving multiple factors, including hypertension, diabetes, chronic tubulointerstitial disease, and other metabolic disorders[2]. These pathogenic factors not only directly affect kidney structure but also interact through various pathways, ultimately exacerbating the decline in kidney function. Hypertension is a critical risk factor for CKD, as it elevates renal blood pressure and impairs glomerular filtration, thereby accelerating renal disease progression.[3]. Diabetes, particularly diabetic nephropathy, has become one of the leading causes of CKD. Hyperglycemia can induce metabolic imbalances, oxidative stress, and inflammatory responses, resulting in damage to both the renal tubules and glomeruli[4]. Furthermore, Acute Kidney Injury is closely associated with the onset of CKD. Studies have shown that individuals who have experienced AKI are at significantly higher risk of developing CKD later, suggesting that AKI may serve as an important early warning signal for CKD progression[5]. Although the mechanisms of CKD progression are complex, early identification and intervention remain key to improving patient outcomes. The subtle early symptoms of CKD complicate timely diagnosis, highlighting the critical need for novel biomarkers to enable early detection and intervention. This is essential for reducing CKD incidence and improving patient outcomes.

In recent years, inflammatory responses have played a key role in the onset and progression of CKD. C-reactive protein (CRP), a key acute-phase reactant, is markedly elevated during inflammation and is commonly used to assess inflammatory states in various diseases[6]. Concurrently, albumin (ALB), a primary indicator of nutritional status and liver function, holds significant diagnostic value due to its altered levels in a range of pathological conditions. Hypoalbuminemia is common in CKD patients and is often associated with further renal function decline and poor prognosis[7]. The CAR, as an emerging composite marker, has garnered widespread attention in recent years. CAR combines the assessment of inflammation and nutritional status, offering significant prognostic value in a range of clinical conditions, including cardiovascular diseases, metabolic disorders, and cancers[8–10]. However, the application of CAR in CKD remains in the exploratory phase, with most studies focusing on CRP or ALB individually, rather than on a comprehensive evaluation of both markers. Investigating CAR’s potential role in CKD is clinically critical for enhancing early diagnosis and predicting treatment outcomes.

This study aims to explore the relationship between the CAR and CKD by analyzing data collected from the National Health and Nutrition Examination Survey (NHANES) database between 1999 and 2010. This study aims to assess the distribution of CAR values and CKD prevalence within the population, providing novel insights for early diagnosis and risk prediction, while offering a theoretical foundation for the development of clinical intervention strategies.

## 2 Materials and Methods

### 2.1 Study Population

The data for this study were obtained from the NHANES database, covering six survey cycles from 1999 to 2010. Basic demographic information, lifestyle factors, and biochemical indicators of the adult population in the United States were collected through household interviews, mobile examination centers, and laboratory tests. This database is publicly available, and researchers have free access to it. The study procedures were approved by the National Center for Health Statistics Institutional Review Board, and all participants provided written informed consent. To ensure participant privacy, all personal identifying information was de-identified during the research process.

During the data processing phase, we conducted screening based on the research objectives, excluding 26,781 individuals under the age of 18, 4,083 individuals lacking data on CKD or the CAR, and 1,247 pregnant women. Ultimately, 30,049 eligible participants were included in the analysis to assess the correlation between CAR and CKD ( Figure 1).

**Fig. 1.**
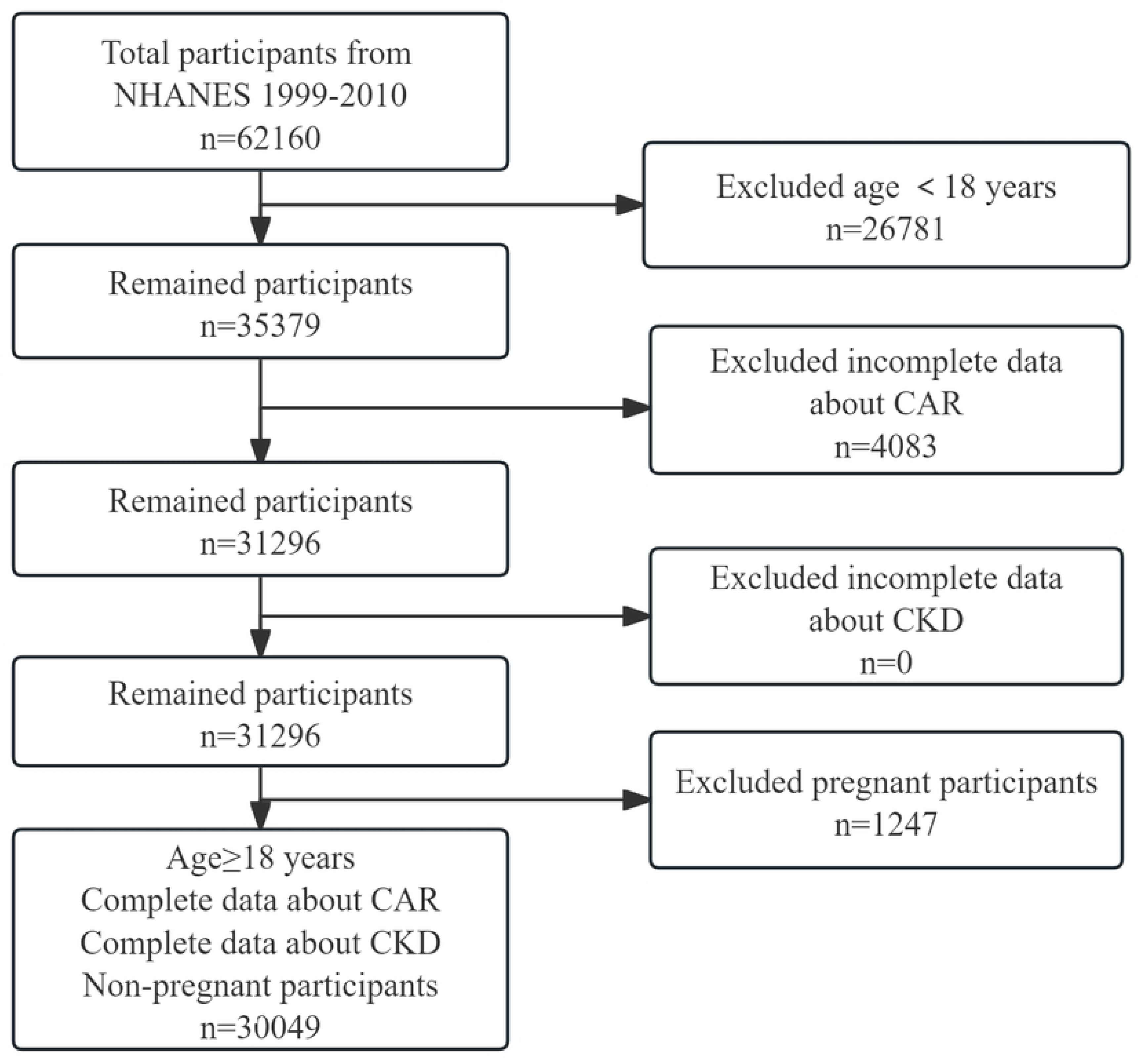
Flow chart of sample selection from the 1999-2010.

### 2.2 Research Variables

#### 2.2.1 Definition of CKD

In this study, CKD was determined based on the definition provided by the Kidney Disease: Improving Global Outcomes Chronic Kidney Disease Work Group[4]. The diagnosis of CKD includes one or more of the following criteria, with symptoms persisting for three months or longer: an estimated glomerular filtration rate below 60 mL/min/1.73 m², calculated using the CKD Epidemiology Collaboration serum creatinine equation; or a urinary albumin-to-creatinine ratio greater than 30 mg/g.

#### 2.2.2 CAR

In this study, the exposure variable, CAR, was calculated as the ratio of CRP (mg/L) to ALB (g/L). Measurements of CRP and ALB were conducted at the NHANES mobile examination centers and were subsequently sent to designated laboratories for further analysis. All blood samples were rigorously screened prior to collection to ensure they met a set of exclusion criteria, ensuring the validity and reliability of the data. To guarantee accuracy, NHANES implemented strict quality control measures throughout the sample collection and analysis process, ensuring high-quality and consistent measurement data.

#### 2.2.3 Study Covariates

Based on existing literature and clinical relevance, this study collected multiple covariates potentially associated with CKD[11, 12]. Data were obtained from the Centers for Disease Control and Prevention through a computer-assisted personal interview system, covering participants’ demographic characteristics, lifestyle, health status, physical measurements, and biochemical indicators.

##### 2.2.3.1 Demographic Variables

Key demographic variables include age, gender, race (categorized as non-Hispanic White, non-Hispanic Black, Hispanic, Mexican American, and other), education level (high school or lower, high school graduate, and post-high school), marital status (unmarried, married, or cohabitating), and poverty income ratio (PIR, classified as <1.3, 1.3–3.5, and >3.5).

##### 2.2.3.2 Lifestyle Variables

Lifestyle-related variables include smoking status and alcohol consumption. Smoking status is classified as: never smoked (smoked fewer than 100 cigarettes in total), former smoker (has smoked but does not currently smoke), and current smoker (currently smoking). Alcohol consumption is categorized as: regular drinker (drinks at least 12 alcoholic beverages per year) and non-drinker (drinks fewer than 12 alcoholic beverages per year).

##### 2.2.3.3 Health Status Variables

Self-reported health status variables include cancer, diabetes, hypertension, and cardiovascular disease(CVD). The diagnosis of diabetes includes a doctor’s confirmation of diabetes, current use of diabetes medication, fasting plasma glucose > 7.0 mmol/L, or glycated hemoglobin (HbA1c) ≥ 6.5%. Hypertension is diagnosed based on one or more of the following criteria: systolic blood pressure ≥ 140 mmHg, diastolic blood pressure ≥ 90 mmHg, the use of antihypertensive medications, or a self-reported history of the condition. CVD is identified through a medical conditions questionnaire, including whether the participant has been diagnosed with coronary artery disease, congestive heart failure, or heart attack.

##### 2.2.3.4 Physical Measurements and Biochemical Tests

Physical measurement variables include body mass index (BMI), categorized into three groups: >25, 25-30, and ≤ 30 kg/m². Biochemical test indicators include triglycerides (TG), high-density lipoprotein cholesterol (HDL), low-density lipoprotein cholesterol (LDL), alanine aminotransferase (ALT), and aspartate aminotransferase (AST).

### 2.3 Statistical Analysis

All statistical analyses were performed following the recommended standards of NHANES. Continuous variables were reported as means with standard deviations (SD) or interquartile ranges (IQR), while categorical variables were presented as frequencies and percentages. To evaluate baseline differences across CAR quartiles, independent samples t-tests were applied to continuous variables, and chi-square tests were used for categorical variables. Subsequently, restricted cubic splines (RCS) were employed to further investigate the dose-response relationship between CAR and CKD. The association between CAR and CKD was evaluated through multivariate logistic regression analyses in several models. Specifically, Model 1 was unadjusted for confounders, Model 2 adjusted for age, gender, and race, and Model 3 further incorporated marital status, education level, PIR, alcohol consumption, smoking status, BMI, hypertension, diabetes, and CVD as potential confounders. Given that the data contained missing values ranging from 0% to 15%, multiple imputation by chained equations using the random forest algorithm was applied to address the missing data. Subgroup analyses were performed to examine the association between CAR and CKD, stratifying by key factors such as gender, race, hypertension, diabetes, CVD, and BMI. To assess heterogeneity between different subgroups, interaction analyses were also performed. All data processing and statistical analyses were conducted using R software (version 4.4.0), with statistical significance set at a P-value of <0.05.

## 3 Results

### 3.1 Baseline Characteristics of Participants

A total of 30,049 participants were included in this study, of whom 24,626 (81.95%) had no CKD and 5,423 (18.05%) were CKD patients. The baseline characteristics are shown in Table 1. The results demonstrated a significantly elevated C-reactive protein to albumin ratio (CAR) in the CKD group compared to the non-CKD group (0.15 vs 0.09, P<0.01). In terms of lipid profiles, the CKD group had higher levels of triglycerides and LDL (P<0.01), but the difference in HDL was not statistically significant (P=0.932). Additionally, the average age of CKD patients was significantly higher than that of non-CKD participants (59.81 vs. 43.14, P<0.01). Significant differences were identified between the two groups regarding gender, race, education level, marital status, poverty index, BMI, alcohol consumption, hypertension, diabetes, CVD, cancer history, and smoking status (all P<0.01). A significant association was found between CAR quartiles and CKD status (χ²=416.13, P<0.01). These findings indicate a strong association between CAR levels and the onset of CKD, with multiple sociodemographic and lifestyle factors potentially influencing its development and progression.

**Table 1.**
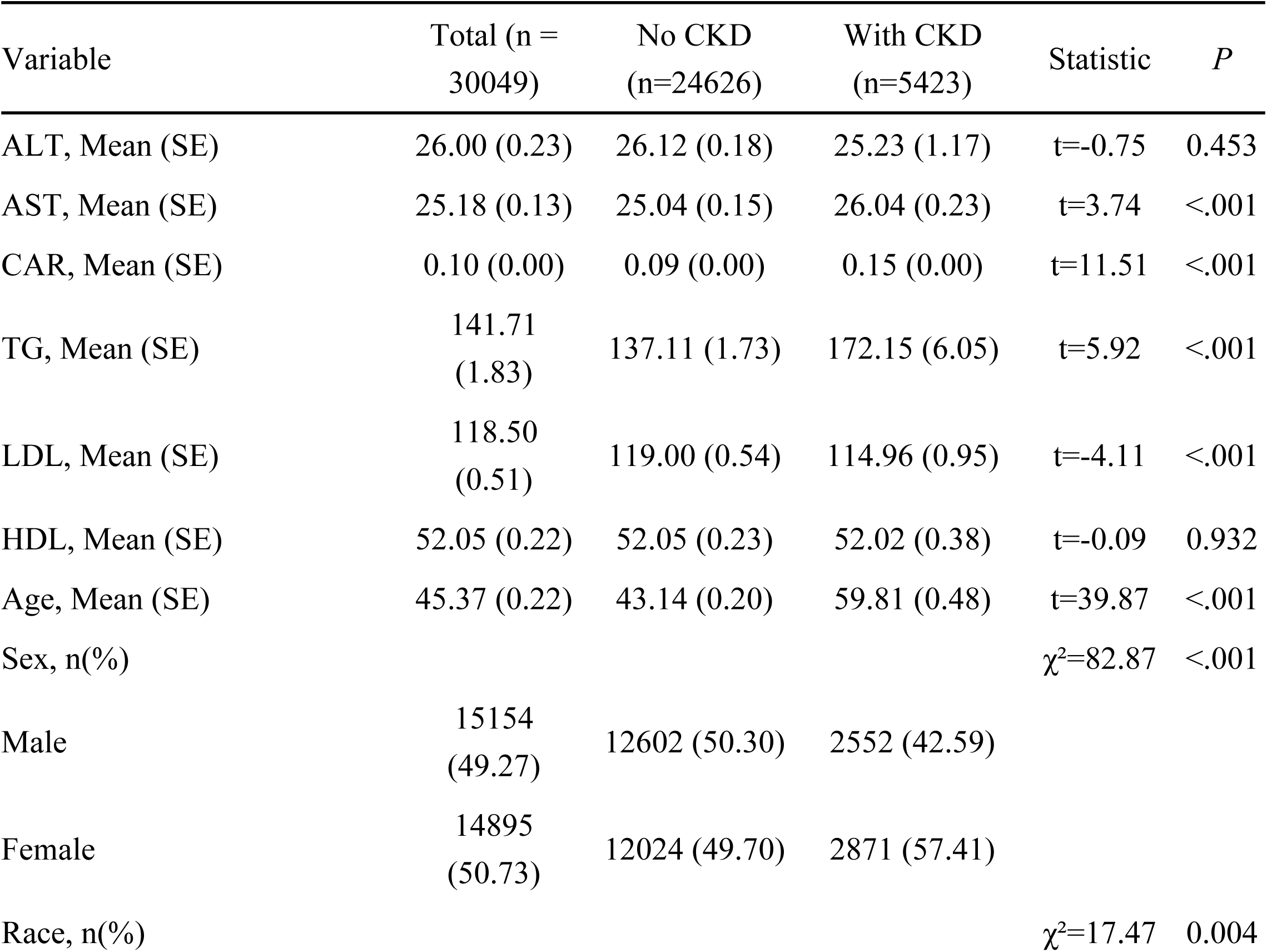
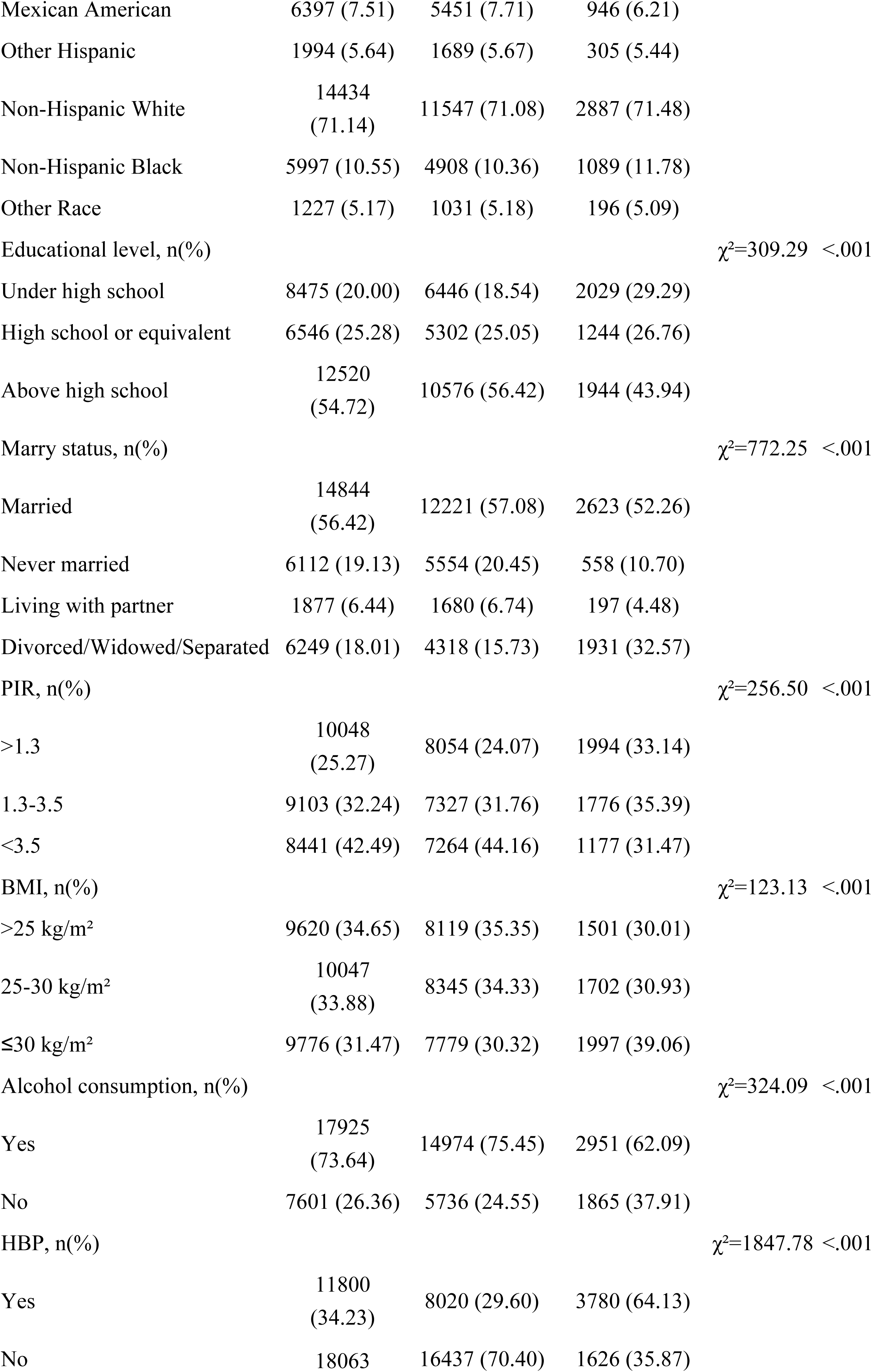
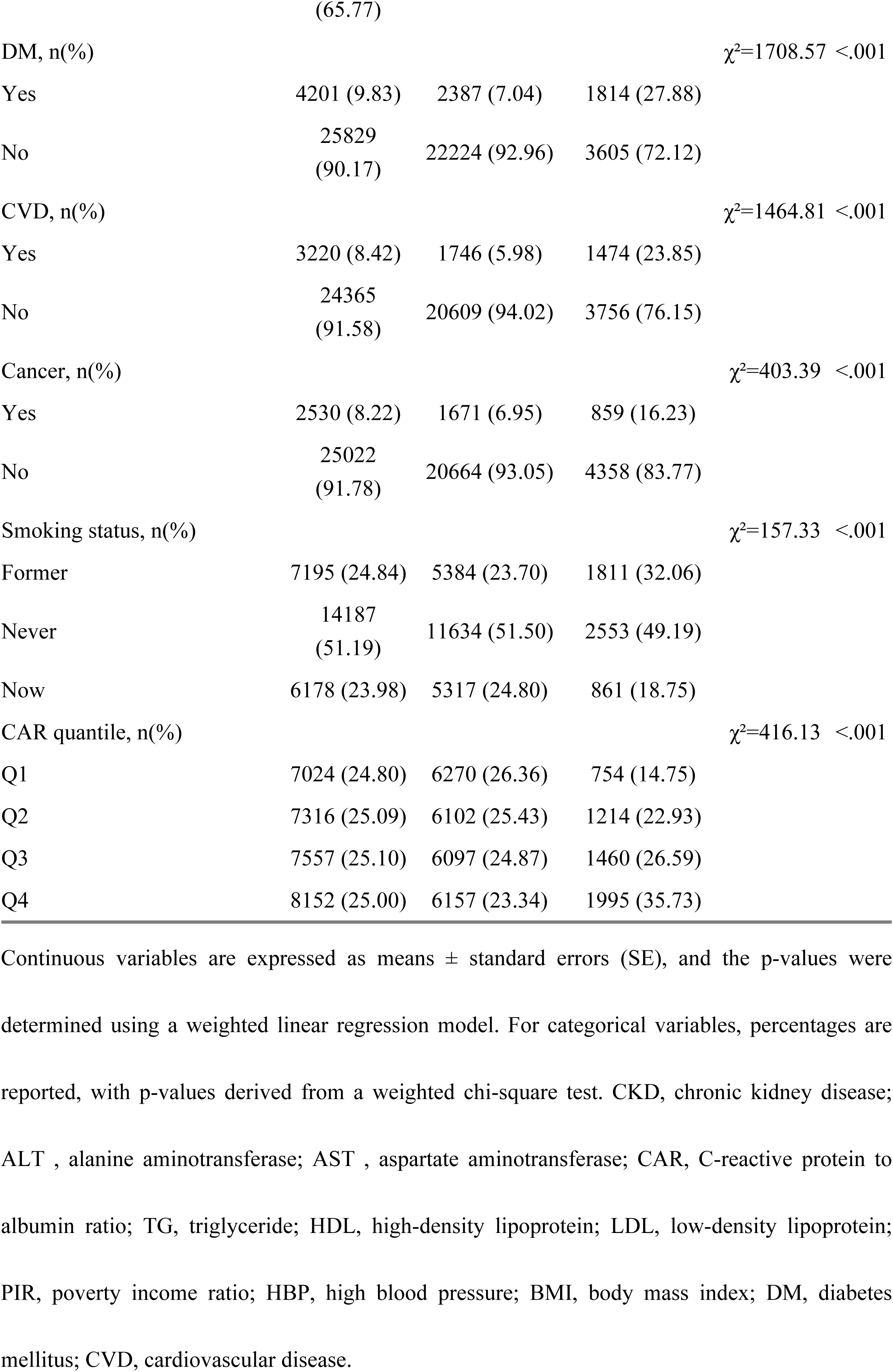
Baseline characteristics of the study cohort.

### 3.2 Association Between CAR and CKD

To further investigate the relationship between CAR and CKD using the NHANES database, a multivariable logistic regression model was applied to evaluate the association between CAR levels and CKD risk. As shown in Table 2, in the unadjusted baseline model, CAR levels were strongly positively correlated with CKD risk (OR = 2.59, 95% CI = 2.19 ∼ 3.07, P < 0.01). After adjusting for basic demographic variables such as gender, age, and race (Model 2), the association between CAR and CKD was somewhat weakened but still significant (OR = 1.98, 95% CI = 1.70 ∼ 2.31, P < 0.01). When additional socioeconomic and lifestyle factors were included (Model 3), the positive correlation between CAR and CKD remained evident and significant (OR = 1.76, 95% CI = 1.34 ∼ 2.31, P < 0.01). When CAR was analyzed as a categorical variable, the association with CKD risk mirrored the findings from the continuous analysis. Specifically, individuals in the Q2, Q3, and Q4 groups exhibited a significantly higher risk of CKD compared to the Q1 group (lowest ln-transformed CAR), with all comparisons yielding P < 0.05. After adjusting for Model 2, the CKD prevalence in the Q3 and Q4 groups was 1.23 times and 1.71 times higher, respectively, compared to the Q1 group (P < 0.05). After full adjustment, the OR for the Q2, Q3, and Q4 groups compared to the Q1 group were 1.28 (95% CI: 1.03, 1.60), 1.29 (95% CI: 1.05, 1.58), and 1.63 (95% CI: 1.27, 2.09), respectively (all P < 0.05). Furthermore, in all models, the incidence of CKD increased progressively with higher ln-transformed CAR values (P < 0.05), further confirming the robust association between CAR levels and CKD risk.

**Table 2.** The relationship between CAR and CKD.

| Variables | Model1 |  | Model2 |  | Model3 |  |
| --- | --- | --- | --- | --- | --- | --- |
|  | OR<br>(95%CI) | P | OR<br>(95%CI) | P | OR<br>(95%CI) | P |
| CAR | 2.59 (2.19 ~ 3.07) | <.001 | 1.98 (1.70 ~ 2.31) | <.001 | 1.76 (1.34 ~ 2.31) | <.001 |
| CAR quantile |  |  |  |  |  |  |
| 1 | 1.00<br>(Reference ) |  | 1.00<br>(Reference ) |  | 1.00<br>(Reference ) |  |
| 2 | 1.61 (1.35 ~ 1.93) | <.001 | 1.17 (0.97 ~ 1.40) | 0.105 | 1.28 (1.03 ~ 1.60) | 0.032 |
| 3 | 1.91 (1.63 ~ 2.23) | <.001 | 1.23 (1.05 ~ 1.45) | 0.013 | 1.29 (1.05 ~ 1.58) | 0.020 |
| 4 | 2.73 (2.34 ~ 3.20) | <.001 | 1.71 (1.44 ~ 2.02) | <.001 | 1.63 (1.27 ~ 2.09) | <.001 |

CAR: C-reactive protein to albumin ratio; 95% CI: 95% confidence interval; OR: odds ratio.
Model 1: Crude. Model 2: Adjusted for Age, Sex, and Race. Model 3: Adjusted for age, sex, race,
marry status, educational level, PIR, BMI, smoking status, alcohol consumption, HBP, DM, CVD,
cancer, AST, TG, LDL.

**Fig. 2.**
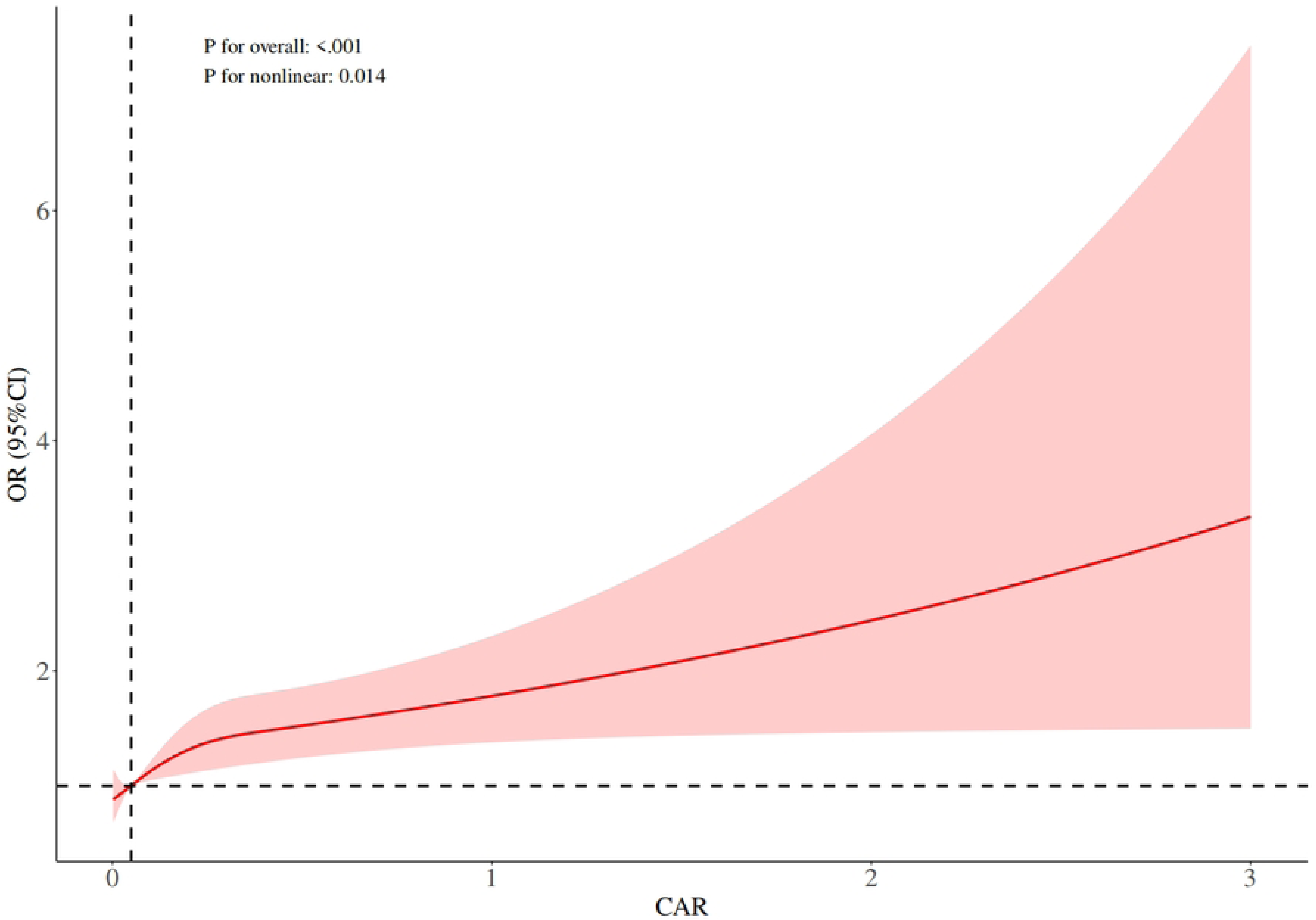
The dose–response relationship between the CAR and CKD was assessed using restricted cubic spline regression, with adjustments for covariates as specified in model 3. The red line represents the odds ratio, and the shaded pink area indicates the 95% confidence interval.

### 3.3 Dose-Response Relationship Between CAR and CKD

Through RCS analysis, we found a significant statistical association between CAR and CKD (overall P-value < 0.01). Further analysis revealed that the relationship between CAR and CKD is nonlinear (nonlinear P-value = 0.014). As CAR values increase, the odds ratio for CKD progressively escalates, highlighting a significant positive correlation between CAR and CKD, characterized by a distinct nonlinear association.

### 3.4 Association Between CAR and CKD: Subgroup and Interaction Analyses

Subgroup analyses revealed significant associations between CAR and CKD risk across various factors, including age, gender, alcohol consumption, hypertension, diabetes, PIR, and smoking status (all P < 0.05), as illustrated in Figure 3. Notably, individuals with at least a high school education, non-Hispanic Whites, non-Hispanic Blacks, those with a BMI ≤30 kg/m² or >25 kg/m², and married individuals exhibited a higher CKD risk associated with CAR. Furthermore, interaction analysis identified a significant interplay between CAR and CKD risk in relation to BMI and education level (interaction P < 0.05). These findings suggest that these factors may be key drivers in the relationship between CAR and CKD risk.

**Fig. 3.**
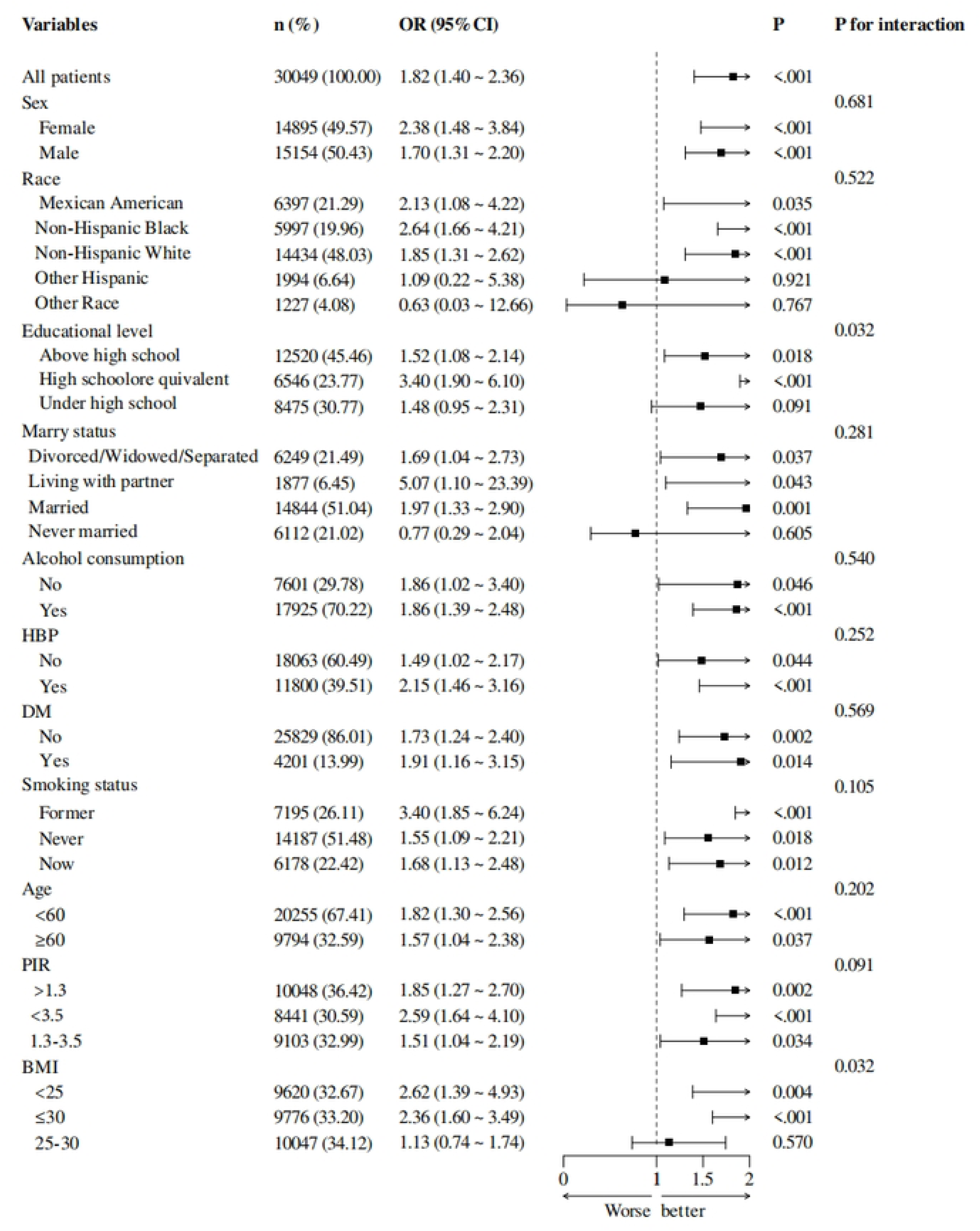
Relationship between NPAR and CKD across subgroup and interaction analyses. Covariates in the multivariable logistic regression models were controlled for as outlined in model 3 of earlier analyses, excluding subgroup-specific variables.

## 4 Discussion

To our knowledge, this is the first study to independently evaluate the relationship between CAR and CKD. The analysis included 30,049 eligible participants, of whom 5,423 were diagnosed with CKD. Multivariable-adjusted models revealed a significant positive correlation between CAR and CKD, with CKD prevalence increasing as CAR levels rose. This finding is biologically plausible, as numerous studies have established a robust link between chronic inflammation and impaired kidney function. Furthermore, as CKD progresses, CAR levels tend to rise, which is consistent with the exacerbation of inflammatory responses and worsening kidney damage. Further analysis suggests that BMI and education level may play an important role in the association between CAR and CKD.

In clinical research, an elevated CAR is frequently correlated with disease progression, organ dysfunction, and adverse outcomes. Notably, previous studies have identified CAR as a key prognostic marker, especially in CVD patients, where an increased CAR is strongly associated with poorer prognosis. This association likely reflects underlying shifts in systemic inflammation and nutritional status, both of which are pivotal factors in CVD management[13]. Studies in CKD patients have identified a significant association between elevated CAR levels and an increased risk of CVD and kidney failure, highlighting CAR as a key predictor of adverse outcomes in this population[14–17]. Chronic inflammation is one of the known pathological mechanisms of CKD and can accelerate the deterioration of kidney function [18, 19]. The CAR serves as an indicator of systemic inflammation, with elevated CAR levels correlating with intensified inflammatory processes, which are a critical driver of CKD progression[20, 21]. As a composite biomarker, the CAR integrates the levels of albumin and CRP, providing an effective measure of systemic inflammation. CRP, an acute-phase reactant predominantly produced by the liver in response to inflammation or infection, is a well-established clinical indicator of inflammatory activity. Research has demonstrated a significant correlation between CRP levels and chronic CKD, particularly in the disease’s onset, progression, and complications, where CRP plays a pivotal role[22–24]. Albumin, a key plasma protein, plays a critical role in maintaining colloid osmotic pressure and facilitating substance transport, making it a crucial marker for assessing nutritional status. It is typically inversely correlated with inflammation, with albumin levels decreasing during chronic inflammatory states, a phenomenon particularly pronounced in CKD patients[25, 26]. Therefore, we hypothesize that elevated CAR is strongly associated with the incidence of CKD and may indirectly influence its onset and progression by reflecting alterations in inflammatory and nutritional status.

The progression of CKD is closely associated with chronic low-grade systemic inflammation. This persistent inflammatory response leads to gradual kidney damage through the activation of the immune system and the secretion of cytokines [27]. Previous studies have demonstrated that inflammation is a key driver of multi-organ pathology in both acute and chronic diseases, with a particularly significant impact on renal pathology[28]. In the onset and progression of CKD, inflammation serves not only as a marker of pathological changes but also as a critical driver of kidney failure[29]. The mechanisms of chronic low-grade inflammation are complex, involving the activation of various cytokines and signaling pathways. For example, the persistent activation of c-Jun NH2-terminal kinase in renal tubular cells is closely related to inflammation and fibrosis in the renal interstitium. The activation of the c-Jun NH2-terminal kinase pathway promotes the expression of fibrogenic factors such as TGF-β1 and CTGF[30]. Moreover, renal resident epithelial cells are crucial in detecting, regulating, and repairing inflammation, with prolonged KIM-1 expression potentially influencing the progression of renal inflammation and fibrosis[31]. Vitamin D deficiency is common in CKD patients and is closely related to the body’s inflammatory and immune responses. Adequate vitamin D levels help reduce the incidence and mortality of CKD patients[32]. At the same time, the oxidative stress and chronic inflammation in CKD patients are closely associated with the development of CVDs, which further aggravate kidney damage and accelerate disease progression[33]. Throughout CKD progression, renal inflammation extends beyond cytokine release to include the activation of multiple signaling pathways, such as NF-κB and Nrf2, which are pivotal in the damage response of renal tubular and glomerular cells. Chronic inflammation contributes to a decline in glomerular filtration rate, potentially progressing to end-stage renal disease[34]. Additionally, studies have demonstrated that renal inflammation is linked to subclinical injury that persists following acute kidney injury, thereby elevating the risk of CKD progression[35].

CAR has emerged as a key biomarker for assessing systemic inflammation, particularly in CKD patients, where it holds significant clinical value. Elevated CAR levels reflect an intensified inflammatory response, often linked to underlying mechanisms such as oxidative stress and immune dysfunction. Given the inherent association between CKD and systemic inflammation, CAR serves as a quantitative measure of the inflammatory burden. Kwon et al. demonstrated that increased CAR levels correlate with higher mortality in kidney transplant recipients, underscoring the pivotal role of inflammation in CKD prognosis[36]. Similarly, Zhang’s study established a significant association between CAR and incident CKD in the Chinese population, emphasizing the critical role of inflammation in disease progression[37]. Additionally, the close relationship between CAR and inflammation has been further validated by other scholars. The research by Ansar and Ghosh pointed out that CRP, an important marker of acute-phase response, significantly increases in tissue injury or infection, thereby contributing to the elevation of CAR[38]. In CKD patients, elevated CRP levels coupled with decreased albumin levels lead to an increased CAR, reflecting the inflammatory burden and highlighting inflammation’s critical role in renal injury. CAR not only serves as an indicator of inflammation but also predicts the worsening of clinical outcomes. Şimşek et al. demonstrated that CAR can predict acute kidney injury in patients with moderate to severe CKD, further reinforcing the central role of inflammation in renal function decline[39]. Similarly, the research by Sant’Ana et al. found a strong association between elevated CAR and increased mortality risk in hemodialysis patients, underscoring CAR’s prognostic significance in CKD management[40]. Therefore, monitoring CAR not only provides strong evidence for the progression of CKD but also offers potential targets for the development of therapeutic strategies aimed at slowing disease progression and alleviating the inflammatory burden.

This study utilized data from the NHANES database, which, due to its stringent data collection standards and large sample size, offers a robust foundation for examining the relationship between CAR and CKD. Through detailed stratification and subgroup analyses, we thoroughly assessed the association between CAR and CKD, and examined the heterogeneity of this relationship across different population groups. However, there are some limitations to this study. Firstly, as a cross-sectional study, we are unable to establish a causal relationship between CAR and CKD. Although our results show a significant association, confirmation of causality requires further longitudinal studies to elucidate the temporal sequence and underlying mechanisms. Secondly, despite accounting for various potential confounding factors, some factors were not sufficiently measured, such as individual genetic background, lifestyle habits, and environmental factors, which may play an important role in the relationship between CAR and CKD. Moreover, the unequal distribution of socioeconomic status and access to healthcare resources may also affect the generalizability of the findings. Therefore, longitudinal studies are essential for further validating our findings. These studies will help uncover the causal relationship between CAR and CKD and explore how related factors contribute to long-term effects. This is of great significance for improving early warning systems for CKD, refining personalized intervention strategies, and optimizing disease management. Furthermore, longitudinal data will provide deeper insights to develop more predictive models and offer more precise guidance in clinical practice.

## 5 conclusion

In conclusion, the study underscores the potential of CAR as a pivotal biomarker for CKD, offering new avenues for advancing early risk stratification, precision therapies, personalized management, and preventive strategies in CKD. Integrating CAR testing into clinical workflows enables the identification of high-risk individuals, facilitating timely preventive interventions and improving overall patient outcomes. Future research should focus on elucidating the molecular mechanisms linking CAR with CKD and exploring its broader clinical applications.

## Acknowledgements

The authors wish to express their sincere gratitude to the NHANES team, including all staff members, researchers, and study participants, for their valuable contributions to the data collection and analysis process.

## 6 Author Contribution Statement

Quanyuan Huang, Xiao Lu and Yufang Yang were responsible for the study design, formal analysis, Investigation, Software, Visualization and data analysis. Quanyuan Huang, Wuchan Zhu, Yuanhua Liang and Jinglan Chen authored the manuscript. Xiao Lu performed manuscript revisions. All four authors reviewed and approved the final manuscript.

## 7 Data Availability Statement

The data utilized in this study is publicly available and can be accessed through the National Health and Nutrition Examination Survey (NHANES) database. The dataset is accessible at the following link: https://www.cdc.gov/nchs/nhanes/index.htm.

## 8 Funding

This study was funded by the National Natural Science Foundation of China (No. 82360872).

## 9 Declaration of Competing Interests

The authors declare that they have no financial or personal conflicts of interest that could be perceived as influencing the research presented in this manuscript.

## Notes

### Competing Interest Statement

The authors have declared no competing interest.

